# Neural evidence for categorical biases in working memory for location and orientation

**DOI:** 10.1101/2020.12.15.422978

**Authors:** Gi-Yeul Bae

## Abstract

Previous research demonstrated that visual working memory exhibits biases with respect to the categorical structure of the stimulus space. However, a majority of those studies used behavioral measures of working memory, and it is not clear whether the working memory representations per se are influenced by the categorical structure or whether the biases arise in decision or response processes during the report. Here, I applied a multivariate decoding technique to EEG data collected during working memory tasks to determine whether neural activity associated with the working memory representation is categorically biased prior to the report. I found that the decoding of spatial working memory was biased away from the nearest cardinal location, consistent with the biases observed in the behavioral responses. In a follow-up experiment which was designed to prevent the use of a response preparation strategy, I found that the decoding still exhibited categorical biases. Together, these results provide neural evidence that working memory representations themselves are categorically biased, imposing important constraints on the computational models of working memory representations.

## 1. Introduction

Our physical environment consists of features and objects that vary along continuous dimensions, but our visual system tends to discretize them into a set of categories (Harnad, 2003). When discriminating two colors from a continuous color space, for example, performance is better when the two stimuli are from different categories (e.g., red vs. blue) than when they are from the same category (e.g., one shade of red vs. another shade of red), even if the physical distances are held constant (Bornstein & Korda, 1984; Uchikawa & Shinoda, 1996). Similarly, visual search for orientation is highly efficient when the target is categorically distinct from the distractors (Wolfe et al., 1992). It was initially believed that these category effects are limited to simple visual features, but studies have demonstrated that categorical perception is more general and arises in the perception of a variety of different features (Fahrenfort et al., 2012; Huttenlocher et al., 2000; Lutz, 1986).

Recent studies have also found robust category effects in working memory tasks (Bae et al., 2015; Hardman et al., 2017; Pratte et al., 2017). For example, Bae et al. (2015) found that the focal color for a given category is remembered more precisely compare to other exemplars in the same category, and color working memory is biased away from the category boundary. Similar effects have been observed for location and orientation (Bae & Luck, 2019; Crawford et al., 2016; Huttenlocher et al., 2000; Pratte et al., 2017; Wei & Stocker, 2015). A recent study found that incorporating category effects into computational models of working memory led to new insights into the longstanding ‘slots-versus-resources’ debate (Pratte et al., 2017).

Although category effects in working memory studies are both common and theoretically important, the majority of studies rely solely on behavior to assess category effects. Because behavioral outputs reflect the combined impact of working memory representations, decision processes, and response processes, it is not clear whether working memory representations per se are biased by the categorical structure of the stimulus space. In the case of orientation working memory, for example, studies have consistently found that working memory reports are biased away from the nearest cardinal orientation (Bae & Luck, 2019a; Pratte et al., 2017; Wei & Stocker, 2015). This may indicate that the orientation representations themselves are categorically biased, but it is also possible that the orientation memory is unbiased but the bias instead arises in the processes that translate the memory representations into a behavioral response. Thus, showing that the actual content of working memory is categorically biased even before the report would provide much stronger evidence that working memory representations per se are influenced by the categorical structure of the stimulus space.

The recent development of EEG decoding techniques makes it possible to track stimulus-specific information being held in working memory on the basis of the scalp topography of the EEG signal (Grootswagers et al., 2017). The present study attempted to decode EEG signal associated with location and orientation information and sought to find evidence that the neural signal for a given location and orientation are categorically biased. In Experiment 1, EEG was recorded while participants performed a location delayed estimation task in which they remembered one of 16 locations on a given trial and reported the remembered location after a short delay (Figure 1a). Categorical biases in behavior were quantified as the shift in the response distribution relative to the nearest cardinal location. For example, when the stimulus was slightly clockwise relative to the vertical midline, the reported location might be shifted a little farther clockwise. Categorical biases in the EEG signal were quantified as a shift in the distribution of decoded locations. I hypothesized that if the neural signal associated with the specific location is categorically biased, then the decoded location for the stimulus that was slightly clockwise relative to the vertical midline would be shifted a little farther clockwise.

**Figure 1.**
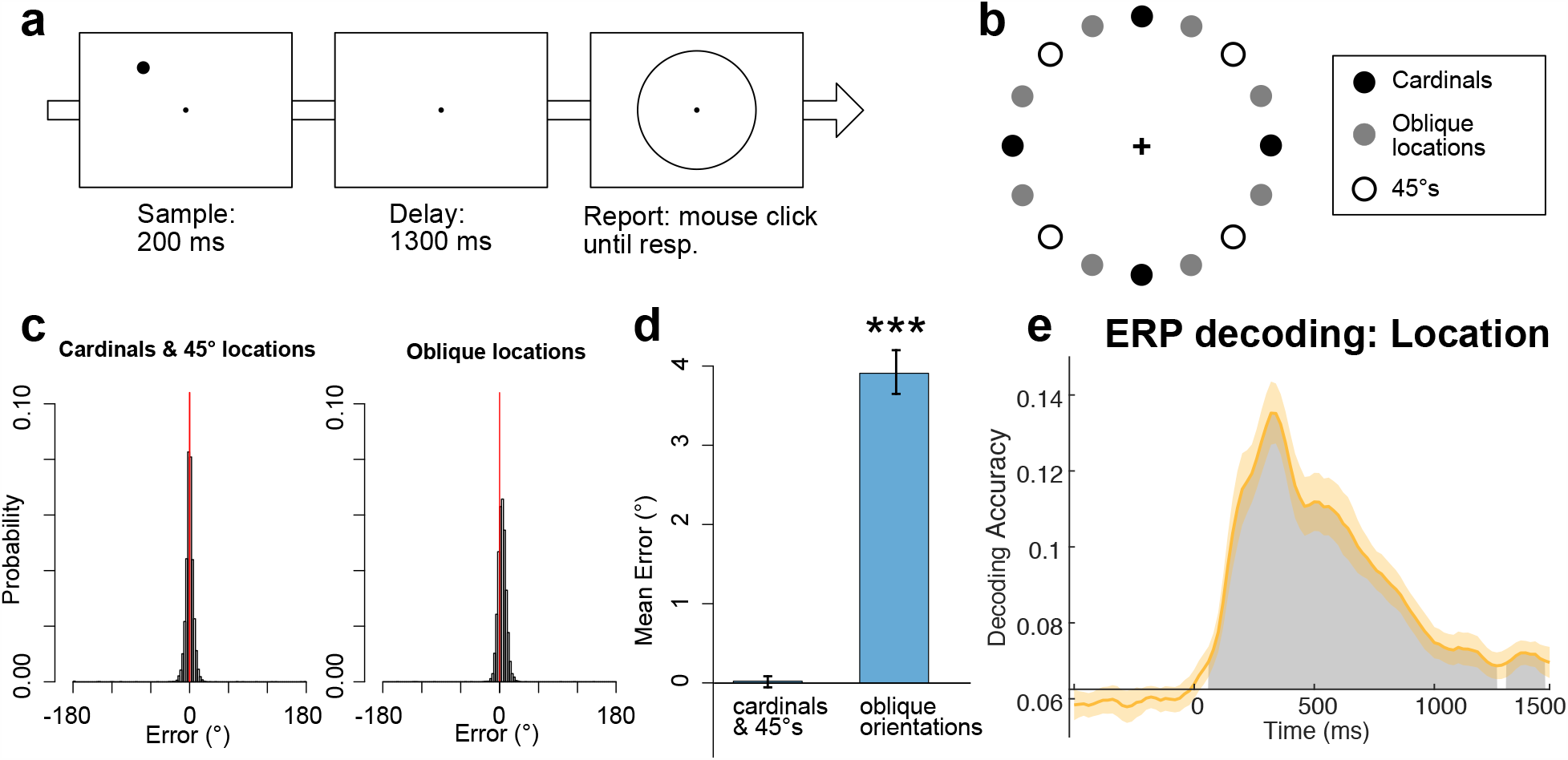
(a) Delayed estimation task used in Experiment 1. On each trial, a stimulus dot was presented and participants clicked the remembered location of the stimulus on the response ring after a 1300-ms delay period. (b) 16 stimulus locations used in Experiment 1. (c) Probability distribution of response errors in the delayed estimation task in Experiment 1, collapsed across all participants, separately for cardinal/45° locations and oblique locations. Positive sign was given to response errors away from the nearest cardinal location and negative sign to response errors toward the nearest cardinal location. The sign was randomly given for the trials with cardinal and 45° locations because the nearest cardinal locations were undefined for them. Red vertical line represents 0° error. (d) Mean signed response error for cardinal/45° locations and oblique locations. Positive error indicates bias away from the nearest cardinal location. Error bars indicate ±1 SEM. (e) Mean ERP decoding accuracy, averaged across participants. The grey area indicates clusters of time points where the decoding was significantly greater than chance (=1/16) after correction for multiple comparisons. The orange shading indicates ±1 SEM. *** = *p* < .001

The spatial working memory task in Experiment 1 could potentially be performed by preparing for a response during the working memory delay because the location of the sample dot was always the location where participants needed to click during the report. To prevent this strategy, Experiment 2 used a 2AFC working memory task in which participants memorized one of 16 orientations indicated by a teardrop object and were presented with two probe objects after a short delay. For the report, they had to press one of two buttons on a response device to indicate which of the two probe items matched the orientation of the original teardrop (Figure 4a). Because the locations of the two probe items varied randomly from trial to trial, and because participants used a response modality that was not correlated with the orientation, they could not prepare the response during the delay period. Categorical biases in behavior were assessed by the difference between trials when the two probes were within the same category (i.e., the same quadrant in the orientation space) versus when they were not. As in Experiment 1, categorical biases in the decoding was assessed on the basis of the distribution of decoding errors.

**Figure 4.**
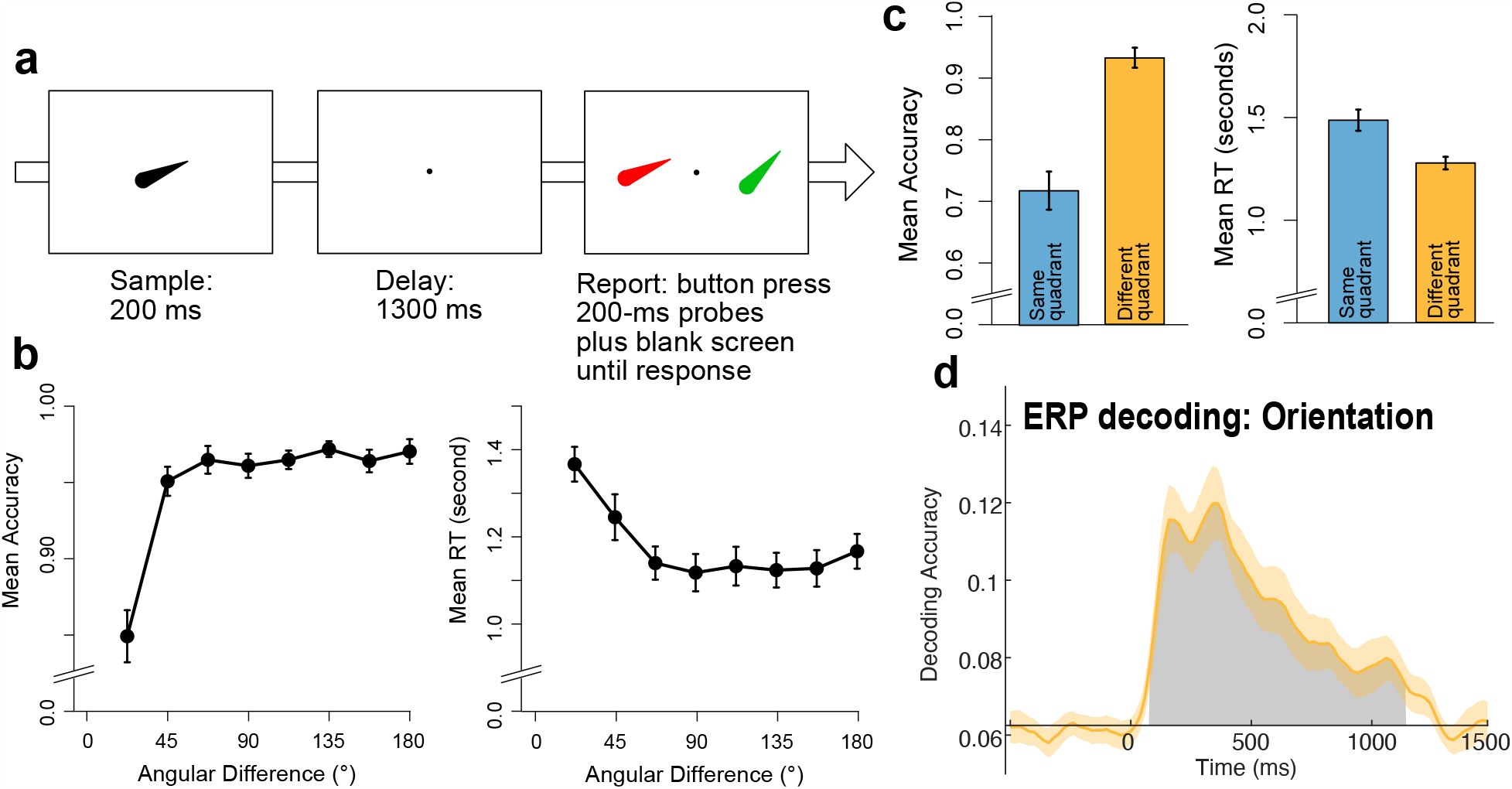
(a) 2AFC task used in Experiment 2. Participants remembered the orientation of the sample teardrop and reported which of the two test teardrops matched the sample teardrop by pressing a button on a gamepad corresponding to the color of the matching teardrop. The positions of the two test teardrops varied unpredictably across trials, but were always centered at mirror-image locations. The test teardrops remained on the screen for 200 ms and a blank screen (not shown in the figure) was presented until response. (b) Mean accuracy and reaction time plotted as a function of the angular difference between the two probe orientations. Error bars indicate ±1 SEM. (c) Mean accuracy and reaction time plotted separately for the trials in which the two probe orientations were within the same quadrant (blue) and for the trials where they were not (orange). The example shown in panel (a) represents a same-quadrant trial (i.e., the two test orientations were in the upper-right quadrant). Error bars indicate ±1 SEM. (d) Mean ERP decoding accuracy averaged across participants. The grey area indicates a cluster of time points where the decoding was significantly greater than chance (=1/16) after correction for multiple comparisons. The orange shading indicates ±1 SEM.

In both experiments, I found strong evidence for categorical biases in both behavioral responses and the EEG decoding. These results demonstrate that the categorical bias in working memory is driven by a bona fide shift of the working memory representations rather than merely by subsequent decision or response properties. These results indicate that computational models of working memory (Bays et al., 2009; Luck & Vogel, 2013; van den Berg & Ma, 2018) must include categorical biases especially when they are compared to alternative models, and they cannot explain away these biases by appealing to decision and response processes.

## 2. Experiment 1: Categorical biases in spatial working memory

### 2.1. Methods and Materials

#### 2.1.1. Participants

Twenty-two college students (16 female, 6 male) between the ages of 18 and 30 with normal or corrected-to-normal visual acuity participated for monetary compensation of $12/hr. The sample size was determined based on related previous studies (Bae & Luck, 2018; Foster et al., 2016). The study was approved by the UC Davis Institutional Review Board and the Arizona State University Institutional Review Board.

#### 2.1.2. Stimuli & Apparatus

Stimuli were generated in Matlab (The Mathworks, Inc.) using PsychToolbox (Brainard, 1997; Pelli, 1997) and were presented on an LCD monitor (Dell U2412M) with a gray background (31.2 cd/m2) at viewing distance of 100 cm. A black fixation dot was continuously present in the center of the display except during the intertrial interval, and participants were instructed to maintain fixation on this dot except during the response period and intertrial interval.

#### 2.1.3. Behavioral Task

Experiment 1 used a standard delayed estimation for spatial working memory (Figure 1a). Each trial started with a 500-ms presentation of the fixation dot followed by a 200-ms presentation of a sample stimulus dot (diameter = .35°). Sixteen discrete locations were used (from 0° to 337.5, in steps of 22.5°, see Figure 1b) and the stimulus dot was presented on an invisible circle with radius of 2.3°. Different locations were tested in random order with equal probability. Participants were instructed to remember the location as precisely as possible over a 1300-ms delay period during which only the fixation dot was visible. A response ring (radius 2.3°) was then presented to indicate that a response should be made. Participants clicked the location on the ring to report the remembered location using a computer mouse. Precision of the report, rather than the speed, was stressed. The next trial started after a 1000-ms delay. Each participant completed a total of 640 trials (40 trials for each of the 16 locations, in random order). Each participant received at least 16 practice trials before beginning the task.

To analyze the behavioral data, I computed response error by taking the difference between the true location and the reported location for a given trial. Critically, I assigned positive sign for the response errors that are in the direction away from the nearest cardinal location (i.e., 0°, 90°, 180°, and 270°) from the target location and negative sign for the response errors that are in the direction toward the nearest cardinal location. Thus, the positive errors indicate that responses were biased away from the nearest cardinal location (i.e., repulsion error) whereas the negative response errors indicate that the responses were biased toward the nearest cardinal location (i.e., attraction error). The positive and negative signs were randomly assigned for cardinal and 45° locations (i.e., 45°, 135°, 225°, and 315°) because the nearest cardinal location is undefined for them.

#### 2.1.4. EEG Recording & Preprocessing

The EEG was recorded using a Brain Products actiCHamp recording system (Brain Products GmbH). Recordings were obtained from 59 scalp sites (FP1/2, AF3/4/7/8, Fz/1/2/3/4/5/6/7/8, FCz/1/2/3/4/5/6, Cz/1/2/3/4/5/6, T7/8, CPz/1/2/3/4/5/6, TP7/8, Pz/1/2/3/4/5/6/7/8/9/10, POz/3/4/7/8, Oz/1/2) for Experiment 1. Electrodes on the left and right mastoids were recorded for use as reference sites. The horizontal electrooculogram (EOG) was recorded from electrodes placed lateral to the external canthi and was used to detect horizontal eye movements; the vertical EOG was recorded from an electrode placed below the right eye and was used to detect eyeblinks and vertical eye movements. Electrode impedances were maintained below 50 KΩ. All signals were recorded single-ended and then referenced offline. The EEG was filtered online with a cascaded integrator-comb antialiasing filter (half-power cutoff at 130 Hz) and digitized at 500 Hz.

Signal processing and analysis was performed in Matlab using EEGLAB Toolbox (Delorme & Makeig, 2004) and ERPLAB Toolbox (Lopez-Calderon & Luck, 2014). The scalp EEG was referenced offline to the average of the left and right mastoids. A bipolar horizontal EOG derivation was computed as the difference between the two horizontal EOG electrodes, and a vertical EOG derivation was computed as the difference between Fp2 and the electrode below the right eye. All the signals were band-pass filtered (non-causal Butterworth impulse response function, half-amplitude cut-offs at 0.1 and 80 Hz, 12 dB/oct roll-off) and resampled at 250 Hz. Portions of EEG containing large muscle artifacts or extreme voltage offsets (identified by visual inspection) were removed. Independent component analysis (ICA) was then performed on the scalp EEG for each subject to identify and remove components that were associated with blinks (Jung et al., 2000) and eye movements (Drisdelle et al., 2017). The ICA-corrected EEG data were segmented for each trial from -500 to +1500 ms relative to the onset of the sample. To verify that eye movements did not impact the decoding results, we also conducted a set of decoding analyses in which trials with eye movements were excluded (see Section 2.1.8).

#### 2.1.5. ERP Decoding Analysis

Past studies have used both sustained ERP activities and oscillatory alpha-power (8-12 Hz) to decode stimulus-specific information held in working memory. They showed that the sustained ERP activities reflect location-independent stimulus information whereas alpha-band activities primarily reflect the locus of spatial attention (Bae & Luck, 2018; Foster et al., 2016). Therefore, the present focused on the ERP activity to decode the location information. However, I also conducted the decoding analysis using the spatial pattern of alpha-power and the results are reported in the Supplementary material.

The segmented EEG was low-pass filtered at 6Hz using EEGLAB *eegfilt()* routine. The use of 6Hz low-pass filter was determined a priori on the basis of previous studies (Bae & Luck, 2018, 2019b) to prevent the influence of alpha-band activity (8-12 Hz) because 1) it mainly reflects the shift of spatial attention (Foster et al., 2016; Rihs et al., 2007), 2) it can masquerade as sustained ERPs under some conditions (Mazaheri & Jensen, 2008; van Dijk et al., 2010), and it is a primary source of trial-to-trial variability in ERPs (Luck, 2014). The filtered data was then resampled at 50Hz to increase the efficiency of the analyses. This procedure produced a 4-dimensional data matrix for each participant, with dimensions of time (100 time points), location (16 different values), trial (40 individual trials for each location), and electrode site (the 59 scalp sites).

To classify the spatial pattern of the ERP signal for a given location into one of the 16 locations, I used the combination of a support vector machine (SVM) and error-correcting output codes (Dietterich & Bakiri, 1995). This model was implemented through the Matlab *fitcecoc()* function. The data were decoded separately for each of the 100 time points from -500 ms to +1480 ms. The decoding for each time point used a 3-fold cross-validation procedure. Trials were divided into three groups and they were averaged together, producing a scalp distribution for the time point being analyzed (a matrix of 3 groups x 16 locations x 59 electrodes). 3-fold cross-validation was chosen a priori based on previous studies (Bae & Luck, 2018, 2019b) to maximize the signal-to-noise ratio for the averaging by increasing the number of trials for each group (Grootswagers et al., 2017). Data from two groups were used to train 16 SVMs (one-versus-all coding) and the remaining data was used to classify the test data into one of 16 locations. The testing was implemented using the Matlab *predict()* function. Decoding accuracy was then computed by comparing the true orientation labels with the classified labels. I also preserved the classified labels from each decoding to analyze which location was more frequently selected for a given test data. This procedure was repeated three times, once with each group of data serving as the test dataset. To minimize idiosyncrasies associated with the assignment of trials to groups, I iterated the entire procedure 10 times with new random assignments of trials to the three groups. The iteration process is necessary to increase the reliability of the decoding and the number of iterations was determined a priori on the basis of previous studies (Bae & Luck, 2018, 2019b).

After completing all the iterations of the cross-validation procedure, decoding accuracy was collapsed across the 16 locations, across the three cross-validations, and across the 10 iterations, producing a decoding percentage for a given time point that was based on 480 decoding attempts (16 locations x 3 cross validations x 10 iterations). After this procedure was applied to each time point, the averaged decoding accuracy values were smoothed across time points to minimize noise using a 5-point moving window (equivalent to a time window of ±40 ms).

In a follow-up analysis, I investigated whether anterior electrode sites and posterior electrode sites exhibit differential decoding and categorical biases (see Section 2.1.7). The decoding procedure of this analysis was identical to the main decoding analysis except for the electrodes used for the decoding (25 anterior electrodes: FP1/2, AF3/4/7/8, Fz/1/2/3/4/5/6/7/8, FCz/1/2/3/4/5/6, Cz/3/4; 25 posterior electrodes: CP3/4/5/6, TP7/8, Pz/1/2/3/4/5/6/7/8/9/10, POz/3/4/7/8, Oz/1/2)(see Figure 3a for the spatial arrangement of electrodes).

**Figure 3.**
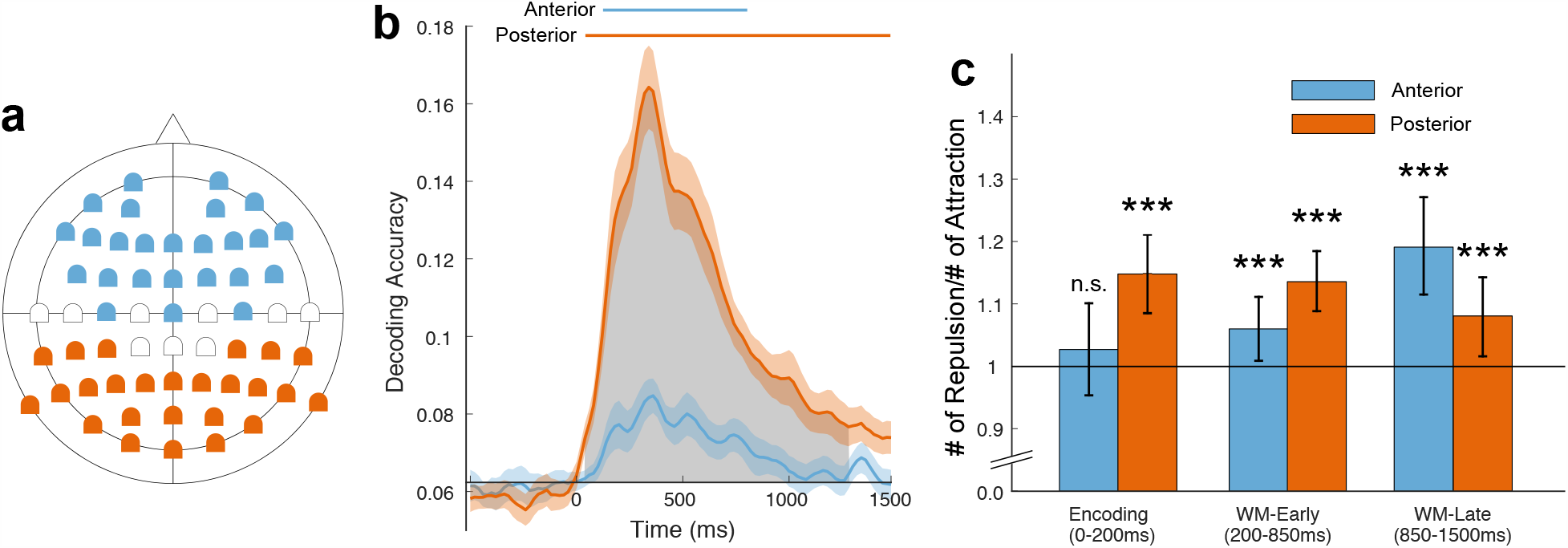
(a) 59 scalp-electrode sites used in Experiment 1. Main decoding analysis were based on the pattern of ERPs over the all the electrodes (i.e., blue, orange, and white). Additional decoding analyses were conducted based on the pattern of ERPs over the anterior electrode (blue) and posterior electrode (orange). (b) Mean ERP decoding accuracy from the decoding with the anterior electrodes (blue) compared to the decoding with posterior electrodes (orange). The blue and orange horizontal bars on top represent significant cluster of time points for the decoding with anterior electrodes and posterior electrodes against chance (1/16). The grey area indicates clusters of time points where the decoding with posterior electrodes was significantly greater than the decoding with anterior electrodes. The shadings along the mean decoding accuracy indicates ±1 SEM. (c) The ratio between the number of repulsion errors and the number attraction errors for the durations of WM encoding (0-200 ms), early WM maintenance (200-850 ms), and late WM maintenance, separately for the decoding with anterior electrodes and posterior electrodes. Horizontal black line represents the hypothetical baseline where the numbers of repulsion and attraction errors are identical (i.e., zero bias). Error bars indicate ±1 SEM. *** = *p* <.001 against chance, two-sided permutation test

#### 2.1.6. Statistical analysis of decoding accuracy

To compare decoding accuracy to chance (= 1/16) at each time point while controlling for multiple comparisons, I used a nonparametric cluster-based permutation analysis (Bae & Luck, 2019d) which is analogous to the cluster-based mass univariate approach that is commonly used in EEG research (Maris and Oostenveld, 2007; Groppe et al., 2011). This method is appropriate here because there was no a priori hypothesis about the time points at which the decoding would be significant and because decoding accuracy may not be normally distributed.

To perform the analysis, I first tested whether the obtained decoding accuracy at each individual time point during the 1500-ms period after the stimulus onset was greater than chance (=1/16) using one-sample t-tests. I then found clusters of contiguous time points for which the single-point *t* tests were significant (*p* < .05), and the *t* scores within each such cluster were then summed together to produce a cluster-level *t* mass. Each cluster-level *t* mass was then compared against an empirical null distribution for the cluster-level *t* mass.

To create an empirical null distribution, I randomly permuted the true labels for the test data prior to computing decoding accuracy (Bae & Luck, 2019d). By permuting the true label for the test data set, the decoding accuracy necessarily at chance. Critically, I used the same permuted label for each time point in a given EEG epoch, to account for temporal autocorrelation in the data during the construction of the empirical null distribution. After computing decoding accuracy with permuted labels, the decoding accuracy was smoothed with a 5-point running average filter (the same procedure applied to the actual decoding accuracy). I then found clusters of contiguous time points using the method described above. If there were no significant clusters of time points, then the cluster mass was set to zero for that permutation trial. If there were more than one cluster of significant time points, I took the mass of the largest cluster.

I repeated the whole permutation process 1000 times with randomly permuted location labels for the test data and constructed the empirical null distribution of the cluster-level *t* mass, resulting in the resolution of *p* = 10^−3^. I then computed the *p* value corresponding to each cluster in the actual data set by examining where each observed *t* mass fell within the null distribution. The *p* value for a given cluster was set based on the nearest percentiles of the null distribution. If the obtained cluster-level *t* mass is larger than the maximum of permuted cluster-level *t* mass, then I reported *p* < .001. I rejected the null hypothesis and concluded that the decoding was above chance (i.e., one-tailed) for any observed cluster-level *t* mass that was in the top 95% of the null distribution (alpha = .05).

To compare decoding accuracy between anterior-based decoding and posterior-based decoding, I created the null distribution by randomly permuting participants between the anterior- and posterior-based decoding 1000 times and used two-tailed test to statistically compare the difference between them.

#### 2.1.7. Analysis of categorical bias in the decoding

Analysis of decoding accuracy focused on the proportion of decoding trials with exact match between the actual label and the classified label. This necessarily overlooks ‘near misses’ which may also contain information about the neural representation for a given stimulus. To investigate how decoding response errors are distributed with respect to the categorical structure of the location space, I assigned positive sign to the decoding response errors that are away from the nearest cardinal location (i.e., repulsion errors) and negative sign to the decoding response errors that are toward the nearest cardinal location (i.e., attraction errors). For the cardinal location themselves (i.e., 0°, 90°, 180°, 270°) and the 45° locations (i.e., 45°, 135°, 225°, 315°), I randomly assigned positive or negative signs to the decoding response error because the nearest cardinal location is undefined for them. Therefore, I focused on oblique locations to investigate the categorical biases.

To quantify categorical biases in the decoding, I compared the frequency of repulsion errors and the frequency of attraction errors by taking the ratio between them (i.e., # of repulsion error divided by # of attraction error). The ratio of one would indicate no categorical biases because it means that there were equal number of repulsion and attraction errors. If, however, the ratio is greater than one, that would indicate that decoding responses were biased away from the nearest cardinal location. Likewise, if the ratio is smaller than one, that would indicate that decoding responses were biased toward the nearest cardinal location. Decoding errors outside the range of ±45° were excluded from this analysis because they are not likely to reflect true location representation given that 99.7% of the behavioral errors were within ±45°.

I computed the ratio for each individual participant for the time points during perceptual encoding period (0-200 ms), for the first half working memory maintenance period (200-850 ms) and the second half of the working memory maintenance period (850-1500 ms). The data within each time period was collapsed to increase the reliability of the bias estimate (i.e. the ratio). Statistical test for the ratio against the null (i.e., the ratio is equal to one) was done via permutation test because the ratio may not be normally distributed. To perform the permutation test, I randomly assigned positive or negative sign to decoding response errors (under the null hypothesis that the decoding errors are independent from the category structure) and computed the ratio. This was repeated 1000 times to construct the null distribution of the ratio (resolution of *p* = 10^−3^). The actual ratio was then compared against the null distribution and I rejected the null hypothesis if the actual ratio was either above the top 97.5% or below 2.5% of the null distribution (alpha = .05, two-tailed).

For the statistical test comparing the ratio between conditions, I randomly permuted participants between the conditions and computed the difference of the ratio from the permuted data and constructed the null distribution of the difference by iterating the process 1000 times. The actual difference between conditions were compared against the null distribution and the null hypothesis was rejected if the actual difference fell either above the top 97.5% or below 2.5% of the null distribution (alpha = .05, two-tailed).

## 3. Experiment 1 Results and Discussion

### 3.1. Experiment 1 Behavior

Figure 1c represents signed response error distribution separately for cardinal/45° locations and oblique locations (see Figure 1b) from Experiment 1. Positive errors indicate that responses were away from the nearest cardinal location and negative errors indicate that responses were toward the nearest cardinal location. The mean of the signed response error for the oblique locations was significantly greater than zero (t(21) = 13.58, *p* < .001, *d* = 2.90), indicating that the reports were biased away from the nearest cardinal location (Figure1d). This result is consistent with previous studies that demonstrated the role categorical structure on perception and working memory (Bae et al., 2015; Huttenlocher et al., 2000).

Although the present study focused on categorical biases, I tested whether cardinal locations (i.e., 0°, 90°, 180°, 270°) exhibited higher precision compared to oblique locations (i.e., all the remaining locations). To that end, I fit a concentration parameter (**κ**) of a von Mises distribution (which has been used to estimate representational precision of a stimulus in a circular space) separately to the response error distribution of cardinal locations and oblique locations. I found that the estimated precision for cardinal locations was significantly greater than oblique locations (cardinals: **κ** = 146.87 vs. obliques: **κ** = 73.05; *t*(21) = 2.18, *p* = .041, *d* = .46), suggesting that the cardinal locations were more precisely reported than the oblique locations.

### 3.2. Experiment 1 Decoding

Figure 1e shows decoding accuracy for Experiment 1. Decoding accuracy began to rise above chance (0.0625 = 1/16) shortly after the onset of the sample stimulus, peaked around 400 ms, and remained high until just before the end of the delay period. The cluster-based permutation test indicated two significant clusters of time points (*p* < .001 for both clusters). Similar results were obtained when the decoding was done after excluding trials with potential eye movements (1 cluster, 80-1500 ms, *p* < .001, one-tailed permutation test)(Supplementary material). These results demonstrate that sustained ERPs contain information about specific location being held in working memory, consistent with previous studies that demonstrated that the spatial pattern of the EEG signal contains location-specific information (Foster et al., 2016).

### 3.3. Experiment 1 Categorical Bias in decoding

Figure 2a shows decoding response error distributions separately for cardinal/45° locations and oblique locations, collapsed across the time points between the stimulus onset and the end of working memory delay (i.e., 0-1500 ms). The bar in the middle of the distribution represents correct decoding (plotted in Figure 1e). More decoding errors occurred near the actual target (i.e., near 0° errors), indicating that the pattern of ERP signals for a given location was more similar to the pattern of ERP signals for similar locations. Importantly, the decoding response error distribution shows more positive decoding errors than negative decoding errors in the distribution for the oblique locations, consistent with the biases in behavior (Figure 1d).

**Figure 2.**
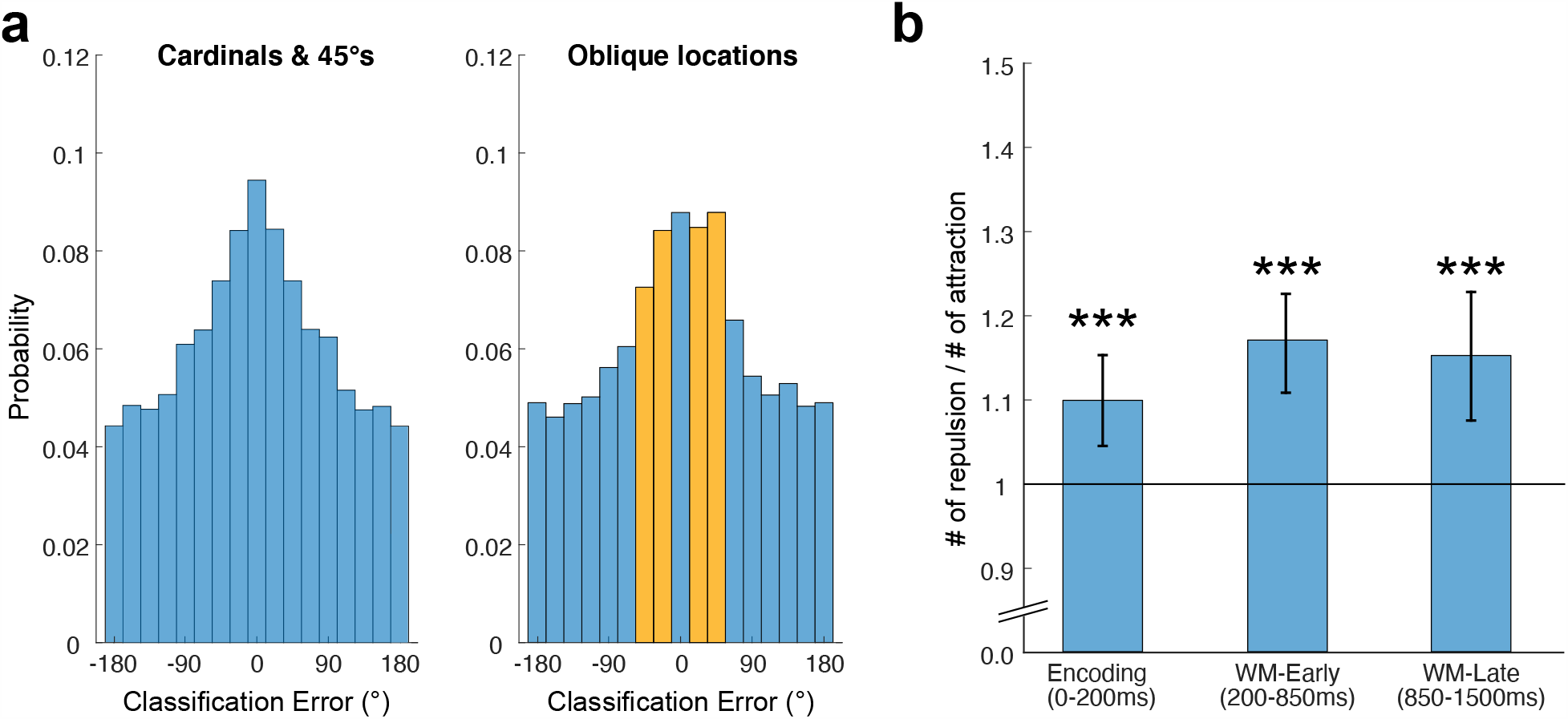
(a) Probability distribution of classification errors for cardinal/45° locations and oblique locations, collapsed across participants and time points (0-1500 ms). Classification errors were computed by taking the difference between the true location label and the predicted label. As in the behavioral data analysis, positive sign was given to the classification errors away from the nearest cardinal location and negative sign to the classification errors toward the nearest cardinal location. The sign was randomly given for cardinal and 45° locations. The middle bar (i.e., 0° classification error) represents decoding accuracy plotted in Figure 1e. (b) The ratio between the number of classification errors away from the nearest cardinal locations (i.e., repulsion) and the number of classification errors toward the nearest cardinal locations (i.e., attraction), plotted separately for the durations of WM encoding (0-200 ms), early WM maintenance (200-850 ms), and late WM maintenance. Classification errors within ±45° range (i.e., orange bars) were used for this analysis because errors outside this range are likely to reflect random noise (see Section 2.1.7). The time points were determined based on the stimulus presentation period (0-200 ms), and the first and the second half of the delay period (200-1500 ms). Horizontal black line represents the hypothetical baseline where the numbers of repulsion and attraction errors are identical (i.e., zero bias). Error bars indicate ±1 SEM. *** = *p* <.001 against chance, two-tailed permutation test.

To quantify this effect, I focused on the decoding response errors within ±45° (i.e., the left two bars and the right two bars from the 0° response error, Figure 2a) and compared the frequency of the positive errors (i.e., repulsion errors) and the frequency of the negative errors (attraction errors) by taking the ratio between them (# of repulsion divided by # of attraction). This was done separately for the time points during the initial stimulus encoding (0-200 ms), the first half of working memory maintenance (200-850 ms), and the second half of working memory maintenance (850-1500 ms) to track the categorical bias over time. As can be seen from Figure 2b, there were more repulsion errors than attraction errors in all three time periods (all *p*s < .001, two-tailed permutation test), supporting the hypothesis that neural representations for spatial location are categorically biased in the same direction as behavior. The same pattern of results was obtained when trials with potential eye movements were excluded prior to the decoding (see Supplementary material), providing evidence that the categorical biases in decoding were not merely driven by systematic eye movements.

Interestingly, the categorical bias during the stimulus encoding seemed weaker than the bias during working memory maintenance. However, they were not statistically different (Encoding vs. Early maintenance: *p* = .183; Encoding vs. Late maintenance: *p* = .257, two-tailed permutation test).

Although the main goal of the present study was investigating categorical biases in the decoding response errors, I attempted to test whether cardinal locations (i.e., 0°, 90°, 180°, 270°) exhibited greater decoding accuracy compared to the decoding accuracy of oblique locations (i.e., all the remaining locations). Result showed no significant difference between them (cardinal: .094 vs. obliques: .090, (t(21) = .80, *p* = .43, *d* = .17).

### 3.4. Categorical biases in anterior and posterior regions

Inspired by recent studies demonstrated that working memory representations in the higher-order processing regions of the brain are more abstract compared to lower-order processing regions (Christophel et al., 2017), I further investigated whether the decoding from anterior and posterior regions would exhibit differential categorical biases. To that end, I conducted two additional decoding analyses: one based on the anterior electrodes (blue electrodes in Figure 3a) and the other based on the posterior electrodes (orange electrodes in Figure 3a). Although the relationship between the underlying neural generator locations and the scalp topography is complex, this posterior-versus-anterior dichotomy served as a coarse proxy for lower-versus-higher levels of processing.

Decoding accuracy for the posterior electrodes was above chance (1 cluster, *p* < .001, one-tailed permutation test) for the most of the time points during the encoding and maintenance periods (40-1480 ms, orange horizontal bar in Figure 3b), and the decoding accuracy for the anterior electrodes was also above chance (1 cluster, *p* < .001, one-tailed permutation test) for much of this period (120-800 ms, blue horizontal bar in Figure 3b). These results are consistent with the hypothesis that the maintenance of stimulus-specific information in working memory is not limited to lower-level visual processing areas but is instead distributed over the cerebral cortex (Christophel et al., 2017). However, the decoding with posterior electrodes sites was much stronger than the decoding with anterior electrode sites (1 cluster, 40-1280 ms, *p* < .001, two-tailed permutation test), suggesting that the information about the specific location stored in working memory is more reliable and stronger in the early visual processing regions.

As in the main decoding analysis, I quantified the categorical bias by taking the ratio between repulsion errors and attraction errors, separately for the anterior and posterior decoding (Figure 3c). Overall, results showed repulsive bias in the decoding error distribution for both anterior-based and posterior-based decoding, but the temporal pattern of the bias was different between them. The bias was not significant during the stimulus encoding period for the decoding with anterior electrodes (*p* = .076) but it became significant during the early maintenance period (*p* < .001) and became even larger during the late maintenance period (*p* < .001). In contrast, the bias of the decoding with posterior electrodes was significantly greater than chance for all three time periods (all *p*s < .001, two-tailed permutation test) with some hint of reduction in the magnitude of the bias as a function of time. To statistically test the differential patterns of the temporal dynamics of categorical bias between the decoding with anterior and posterior electrode sites, I conducted a 2-way ANOVA with time period (encoding, early-WM, late-WM) and electrode region (frontal and posterior) as within-subject factors. I found a significant two-way interaction (*F*(2,42) = 4.158, *p* = .023, η _p_^2^ =.165). Simple main effect analyses showed a significant main effect of time period in the anterior-based decoding (Encoding vs. Early maintenance: *p* = .320; Encoding vs. Late maintenance: *p* = .012, two-tailed permutation test) but no significant main effect of time period in the posterior-based decoding (Encoding vs. Early maintenance: *p* = .432; Encoding vs. Late maintenance: *p* = .676, two-tailed permutation test). These results suggest that higher- and lower-level processing regions exhibit differential temporal dynamics of categorical biases.

## 4. Experiment 2: Categorical biases in orientation working memory

### 4.1. Materials and Methods

#### 4.1.1. Participants

Sixteen college students (10 female, 6 male) between the ages of 18 and 30 with normal or corrected-to-normal visual acuity participated for monetary compensation of $12/hr. They had experience with at least one prior working memory task used in an independent research project. The sample size was determined based on previous studies (Bae & Luck, 2018; Foster et al., 2016). The study was approved by the UC Davis Institutional Review Board and the Arizona State University Institutional Review Board.

#### 4.1.2. Stimuli, Apparatus, & Procedure

Apparatus used in this experiment was identical to Experiment 1. The task in Experiment 2 (Fig. 4a) was designed to prevent the use of the location-based response preparation strategy. On each trial, a tear-drop shaped sample stimulus (2.3° long, 0.7° maximum width) was presented centered on the fixation dot for 200 ms. The orientation of a given teardrop was defined by the angular position of the tip relative to the center of the object. As in Experiment 1, sixteen discrete orientations (from 0°, in steps of 22.5°) were used and the orientation for a given trial was randomly chosen from this set. Participants were instructed to remember the orientation as precisely as possible over a 1300-ms delay period. At the end of the delay interval, two test teardrops appeared for 200 ms, followed by a blank screen (not shown in the figure). One of the two test teardrops matched the orientation of the sample teardrop and one of them differed by some multiple of 22.5° (with an equal likelihood of all multiples between 22.5° and 337.5°). To minimize the shift of covert attention toward the location of test teardrops, one of the test teardrops was presented at random location within a 3° x 3° region centered 4.5° to the left of the fixation dot, and the other appeared at the mirror image position in the right visual field. Each of the test teardrops was presented in a randomly chosen color, selected without replacement from a set of three colors (red, green, and light blue). Each color was associated with one of three buttons positioned on a gamepad, and participants were instructed to press the button corresponding to the color of the teardrop whose orientation matched the orientation of the sample teardrop. The color and location of the matching teardrop was randomized across trials. Because the task did not explicitly require participants to report the location of the matching teardrop and because the location of the test teardrop was randomized across trials, participants were not able to prepare the response. Both accuracy and speed of the response were stressed. After the response, the display blanked, and the next trial started after a 1000-ms delay.

After at least 16 practice trials, each participant completed a total of 640 trials (40 for each of the 16 orientations. Each participant received at least 16 practice trials before beginning the task.

#### 4.1.3. EEG Recording & decoding analysis

EEG recording and preprocessing procedures were identical to Experiment 1 except that recordings were obtained from 27 scalp sites (FP1/2, F3/4/7/8, C3/4, P3/4/5/6/7/8/9/10, PO3/4/7/8, O1/2, Fz, Cz, Pz, POz, and Oz). The decoding analyses, including statistical tests and the analysis of categorical biases, were identical to Experiment 1.

## 5. Experiment 2 Results and Discussion

### 5.1. Experiment 2 Behavior

Figure 4b summarizes the data from the 2AFC task, showing both mean accuracy and mean response time as a function of the orientation difference between the two test teardrops. Accuracy decreased and response time increased when the orientations of the two test teardrops were similar, which reflects the limited precision of the WM representation of the teardrop orientation. To test this pattern statistically, the accuracy and response time data were entered into separate one-way ANOVAs with a factor of orientation difference. The main effect of orientation difference was significant for both mean accuracy (*F*(7,120) = 19.03, *p* < .001, η_p_^2^ =.53 and mean response time (*F*(7,120) = 4.159, *p* < .001, η_p_^2^ =.20).

To test whether the behavior in this task reflects categorical representation of orientation, I focused on the trials in which target orientations were near the cardinal axis (i.e., 22.5°, 67.5°, 112.5°, 157.5°, 202.5°, 247.5°, 292.5°, 337.5°) and were paired with a non-target orientation that differed by 22.5° (i.e., the smallest orientation difference tested) either toward or away from the cardinal orientation. If orientation memory was biased away from the nearest cardinal orientation, then the task should be more difficult when a target orientation was paired with a non-target orientation that is also away from the cardinal orientation (e.g., 22.5° target paired with 45° non-target) compared to when a target orientation was paired with a non-target orientation that is toward the cardinal orientation (e.g., 22.5° target paired with 0° non-target) because the two probe items are in the same quadrant in the former case but not in the latter. In line with this prediction, accuracy was lower (t(15) = -6.64, *p* <.001, *d* = 1.66) and reaction time was slower (t(15) = 5.14, *p* <.001, *d* = 1.29) when the two probe items were in the same quadrant compared to when they were not (Figure 4c). These results suggest that orientation representation in working memory is biased away from the nearest cardinal orientation.

### 5.2. Experiment 2 Decoding

Figure 4d shows decoding accuracy from the 2AFC task. Decoding accuracy rose above chance shortly after the onset of the stimulus and remained above chance almost until the end of the delay interval. Cluster-based permutation test revealed a large cluster of time points that was significantly different from chance (80-1140 ms, *p* <.001). The decoding without trials with potential eye movements also produced a significant cluster of time points (1 cluster, 100-700 ms, *p* <.001, one-tailed permutation test)(see Supplementary material). Consistent with previous studies, this result demonstrates that the specific orientation is decodable on the basis of the spatial pattern of ERP activity and further demonstrate that the specific orientation held in working memory is decodable even when the direct estimation of the remembered orientation is not required and the response cannot be prepared during the delay interval.

### 5.3. Experiment 2 Categorical Bias in decoding

Figure 5a shows the signed decoding response error distributions separately for cardinal/45°orientations and for oblique orientations collapsed across all the time points from 0 ms to 1500 ms. Interestingly, decoding response distributions showed bi-modality with frequent decoding response errors near 180°. This may show that the ERP activity reflects the representation of a line orientation (as opposed to a ray orientation) that is not distinguishable from an orientation 180° away (e.g., orientation in a Gabor patch). However, given that previous studies showed that all 16 different teardrop orientations were decodable based on the ERP signals (Bae & Luck, 2018), it is more likely that the opposite orientation decoding in the present study reflects that participants remembered the orientation of the thick end of the teardrop (as opposed to the tip of the teardrop) on some proportion of the trials.

**Figure 5.**
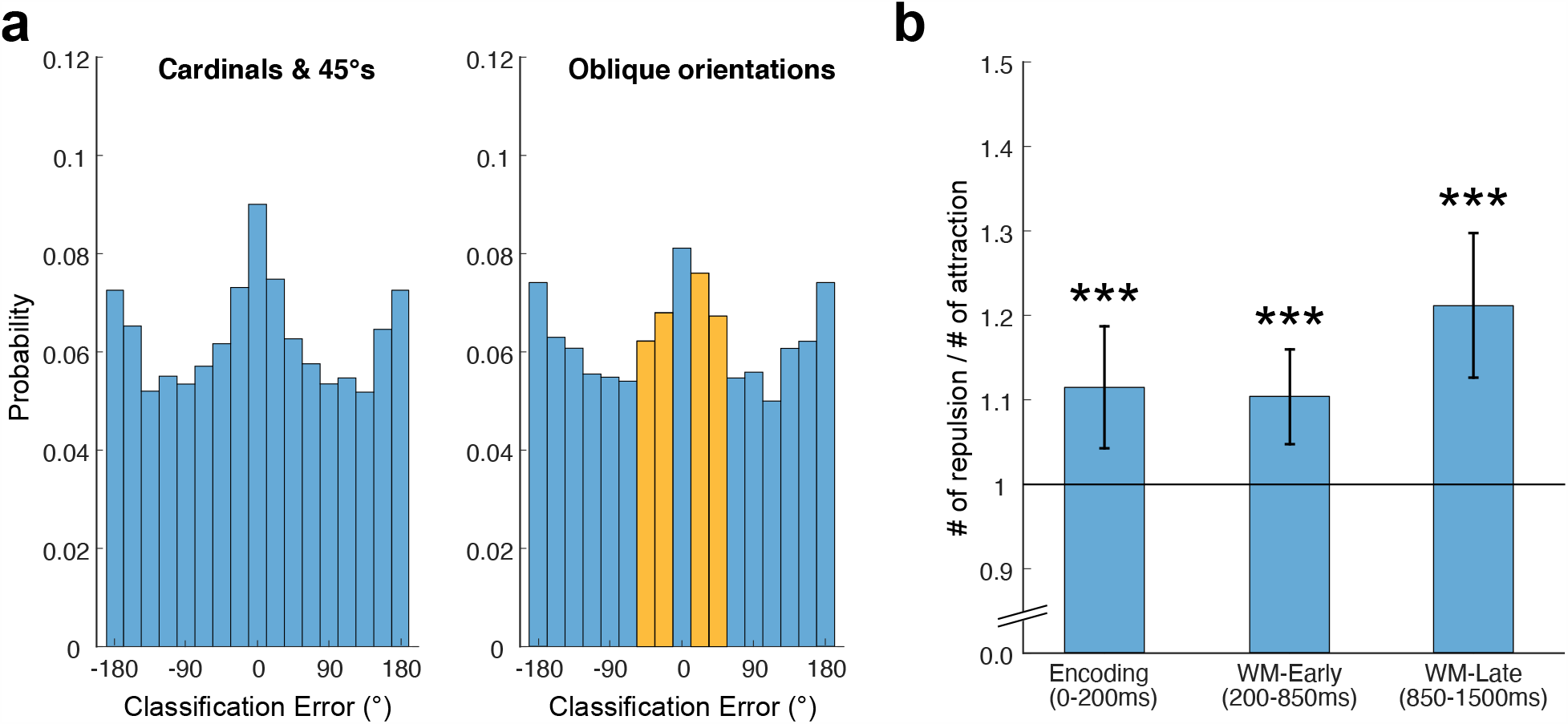
(a) Probability distribution of classification errors for cardinal/45° orientations and oblique orientation in Experiment 2. Greater number of classification errors near ±180° indicates opposite orientation decoding (which may indicate that participants performed that task by remembering the thick-end of the teardrop on some proportion of trials). As in Experiment 1, positive sign was given to repulsion errors and negative sign was given to attraction errors. (b) The ratio between the number of repulsion errors and attraction errors for the durations of WM encoding (0-200 ms), early WM maintenance (200-850 ms), and late WM maintenance. Classification errors within ±45° range (i.e., orange bars) were used for this analysis. Error bars indicate ±1 SEM. *** = *p* < .001

As in Experiment 1, there were more positive decoding response errors (i.e., repulsion errors) than negative decoding response errors (i.e., attraction errors). I compared the frequency of these errors by taking the ratio between them for the duration of stimulus encoding (0-200 ms), early working memory maintenance (200-850 ms), and later working memory maintenance (850-1500 ms). As can be seen from Figure 5b, decoding was biased away from the nearest cardinal orientation starting from the stimulus encoding period, and the bias persisted until the end of working memory maintenance (all *p*s <.001, two-tailed permutation test). The same pattern of results was obtained when trials with potential eye movements were excluded prior to the decoding (see Supplementary material). These results are consistent with the hypothesis that orientation information in working memory is categorically biased before the response is made and provide evidence that the bias is not merely driven by response preparation processes.

The bias seems larger during the late maintenance period, but it was not statistically different from the bias during the early maintenance period (*p* = .164, two-tailed permutation test) and the stimulus encoding period (*p* = .557, two-tailed permutation test).

I also tested whether the cardinal orientations produced greater decoding compared to oblique orientations. Overall, the decoding accuracy for cardinal orientations was greater than the decoding accuracy for oblique orientations (cardinals: .092 vs. obliques: .083), but the difference was not statistically significant (t(15) = 1.48, *p* =. 16, *d* = .37).

## 6. General Discussion

The present study sought to find neural evidence for categorical biases in working memory for location and orientation. Using a multivariate EEG decoding approach, I found that the pattern of neural activity for a location that was near the cardinal axis was more likely to be classified as locations further away from the cardinal axis during both perceptual encoding and working memory maintenance periods. This was consistent with the behavioral response pattern, in which the reports were biased away from the nearest cardinal location. This was replicated in the decoding of orientation in a 2AFC working memory task in which a location-based response preparation strategy was discouraged (if not completely prevented). The 2AFC performance was poorer when the non-target item was within the same category as the target item, and more decoding errors occurred away from the nearest cardinal orientation. These results demonstrate that the categorical structure of the stimulus space—for both location and orientation—biases neural signals associated with working memory representations, suggesting that the categorical bias observed in behavior is driven by the categorically biased representations.

### 6.1. Category effects in working memory

Despite the robust category effects in perception and working memory (Bae et al., 2015; Hardman et al., 2017; Pratte et al., 2017), influential models of working memory do not take the category effects into account when explaining variabilities in working memory behavior (Bays et al., 2009; van den Berg et al., 2012; Luck & Vogel, 2013). Those models typically attribute the variability to stimulus-independent factors such as storage failure (i.e., discrete slot model) or random attentional fluctuations (i.e., continuous resource model). However, a recent study directly demonstrated that the category effects play a critical role in the long-standing ‘discrete slots vs. continuous resources’ debates in working memory (Pratte et al., 2017). When Pratte et al. (2017) incorporated the stimulus-specific variation associated with the category structure of orientation space into working memory models, they found that the discrete slot model provided a better account of visual working memory errors compared to the continuous resource model which otherwise outperformed the discrete slot model. This implies that large amount of variability in working memory errors that was accounted for by the continuous resource model was not random variability but was instead driven by the categorical structure of the stimulus space. Thus, the results from Pratte et al. (2017) further suggest that a failure to account for category-based effects could lead to unwarranted advantage to models that assume random variability in working memory. Here, the present study provides neural evidence that working memory representations per se are categorically biased and further emphasizes that computational models of visual working memory must incorporate the category effect, especially when comparing models for visual working memory.

The present study focused on working memory for location and orientation because the categorical structure for these two features is relatively simple—the category boundaries are well-defined and the size of each category is relatively uniform. However, the lion’s share of visual working research uses color as the memory feature (Allred & Flombaum, 2014) because of the non-spatial nature of the color space. That is, space-based strategies can be completely prevented in color working memory because the location does not provide any meaningful information about what color was presented at that location. However, the categorical structure of the color space is more complex, and it varies depending on the specific color space used in a given study (see Bae et al., 2014, for CIELab vs. HSV color space). In addition, no one has yet successfully decoded color on the basis of the EEG signal (but see, color decoding based on fMRI signal, Brouwer & Heeger, 2009). Thus, it remains to be seen whether the decoding of color would also exhibit categorical biases in a manner that is consistent with behavior.

Research has demonstrated that the categorization also exhibits variabilities in the representational precision in a stimulus-specific way (Bae & Luck, 2019a; Pratte et al., 2017). Experiment 1 of the present study also demonstrated that cardinal locations are more precisely reported than oblique locations. The present study attempted to test whether similar effect occurred in the decoding by comparing decoding accuracy for cardinal locations/orientations and oblique locations/orientations. Although there was some hint of greater decoding accuracy for cardinals compared to obliques, the difference was not statistically significant. However, comparing decoding accuracies is only a weak test for comparing precision differences because decoding accuracy does not fully reflect the noisiness in the neural representations. Quantifying the dispersion of decoding errors (as in behavioral analysis) would be a more appropriate approach, but such approach is not applicable to the present study because the course granularity of decoding response errors (i.e., smallest error = 22.5°) limits accurate estimation of decoding precision. Future study is necessary to investigate whether the categorization process would exhibit differences in the precision of decoding by increasing the number of stimulus in a given stimulus space.

### 6.2. Categorical bias: When and Where

In both experiments, the categorical biases in the decoding occurred during perceptual encoding period and persisted until the working memory maintenance period. This result suggests that the categorical biases were created during the perception of the stimulus and transmitted to the working memory storage. Consistent with this result, a previous study demonstrated that categorical biases occur even when a stimulus was still in view (thus, no explicit memory demand was imposed) using an ‘undelyaed-estimation’ task, supporting the hypothesis that the categorical bias is originated from perception (Bae et al., 2015). However, it is also possible that a working memory process might have taken place during perception, and that process, rather than the perceptual process per se, might have produced the categorical bias. The present study does not provide a clear answer for the exact origin of categorical biases. Future research is necessary to investigate the role of perception and working memory in the categorization process.

Although the location decoding was much stronger when posterior electrodes were used for decoding, the decoding based on anterior electrodes was also significantly greater than chance. Although the relationship between generator sources and scalp sites is complex, this finding suggests that the stimulus-specific information in working memory exists beyond the early sensory regions (Christophel et al., 2017), consistent with previous neuroimaging studies that found content-specific delay activity in frontal areas (Ester et al., 2015). In addition, the present study found that the categorical bias of location memory got stronger with the delay for the decoding with more anterior electrode sites. These results provide new insights on the temporal dynamics of categorical information in the hierarchy of visual processing.

### 6.3. Other biases in working memory

Biases in working memory representations are not only driven by categorical structure of the stimulus space but also by the trial history. Studies on serial dependence have demonstrated that the orientation report on the current trial is biased by the orientation presented in the previous trials (Fischer & Whitney, 2014). Interestingly, recent work suggests that the reactivation of the previous-trial information during the perception of the current-trial stimulus underlies the serial dependence effect (Bae & Luck, 2019b; Barbosa et al., 2020). However, no direct evidence has been provided for which the neural signals for the current-trial stimulus is actually biased in relation to the previous trial stimulus. Thus, finding evidence that the actual representational content of the current trial is biased depending on the stimulus presented in the previous trial would be an important goal of future research.

## Acknowledgement

I thank Steven J. Luck for useful comments on the earlier version of the manuscript, and Aaron Simmons for assistance with data collection.

