## Supplementary material for "Neural evidence for categorical biases in working memory for location and orientation"

**for**

---

---

Gi-Yeul Bae

Department of Psychology
Arizona State University
Tempe, AZ, 85287

**Address for correspondence:**

Gi-Yeul Bae, Ph.D.
Department of Psychology
Arizona State University
950 S. McAllister Ave.
Tempe, AZ 85287
(M) 410-491-5540
(E)

**Contents:**

1. Decoding on the basis of the power of alpha-band (8-12Hz) activity
2. Decoding without trials with potential eye movements
3. Categorical biases in decoding when the decoding was done without trials with potential eye movements

1. <u>Decoding on the basis of the power of alpha-band (8-12Hz) activity</u>

Main decoding analyses focused on the sustained ERP activity associated with the perception and working memory for location (Experiment 1) and orientation (Experiment 2). This focus was based on the previous studies that showed that the sustained ERP activity reflects representational contents held in working memory (Bae & Luck, 2018). However, past studies have shown that the spatial pattern of EEG alpha-band activity can be used to track spatial attention over time in a working memory task (Foster et al., 2016). Here, I attempted to test whether spatial attention is also categorically biased by decoding location (Experiment 1) and orientation (Experiment 2) on the basis of alpha power (8-12 Hz). The decoding procedure was identical to the method reported in the main paper except that the segmented EEG was bandpass filtered at 8-12 Hz (using the EEGLAB *eegfilt()* routine) and the bandpass-filtered EEG was submitted to Hilbert transform to compute the magnitude of the complex analytic signal, and this magnitude was then squared to compute total power in the 8-12 Hz band each time point.

Figure S1a shows the mean decoding accuracy from this analysis. Decoding was above chance for the most of the time points during perception and working memory maintenance. This result is in line with previous study (Foster et al., 2016) and demonstrates that 16 different attended locations can be decoded on the basis of the spatial pattern of alpha power. Figure S1b shows response error distribution from the decoding analysis separately for cardinal/45° locations and oblique locations. As in the main analysis, positive error indicates that the decoding response errors were away from the nearest cardinal location (i.e., repulsion error), and vice versa. Clearly, oblique locations exhibited more frequent repulsion error compared to the attraction errors. When the ratio between them was computed, I found significant repulsion bias starting from perceptual encoding period and the bias persisted until working memory delay period. This result demonstrates that spatial attention is also categorically biased.

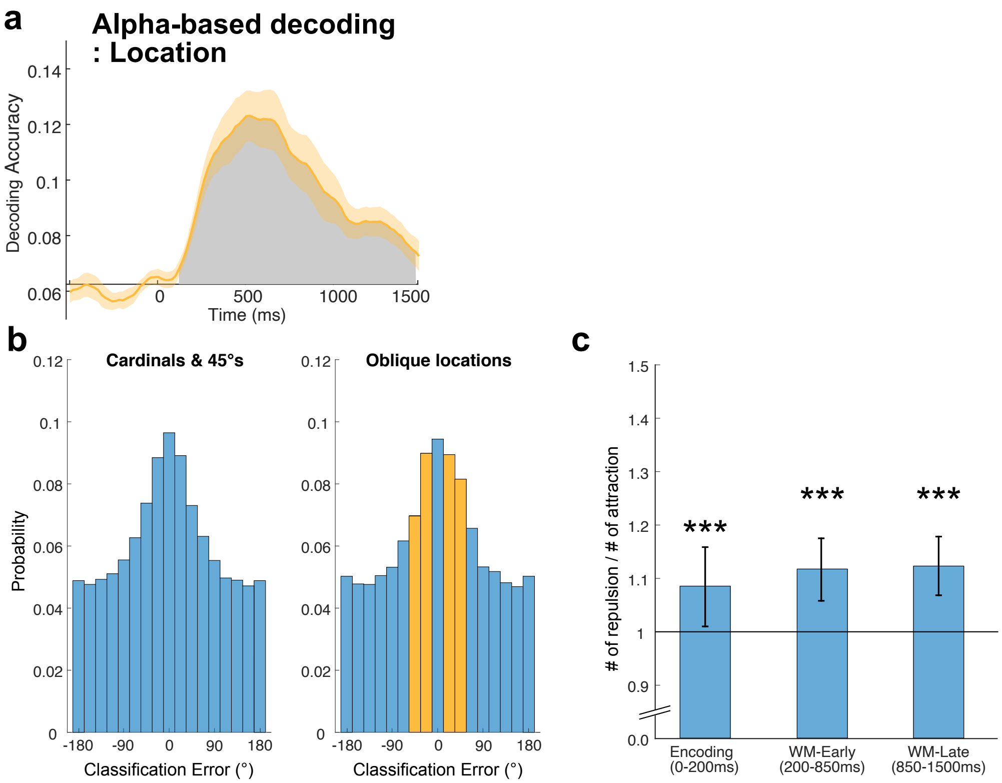


Figure S1. (a) Mean Alpha-based decoding accuracy, averaged across observers in Experiment 1. The grey area indicates clusters of time points where the decoding was significantly greater than chance (=1/16) after correction for multiple comparisons. The orange shading indicates ±1 SEM. (b) Probability distribution of classification errors for cardinal/45° locations and oblique locations, collapsed across observers and time points (0-1500 ms). Classification errors were computed by taking the difference between the true location label and the predicted label. As in the behavioral data analysis, positive sign was given to the classification errors away from the nearest cardinal location and negative sign to the classification errors toward the nearest cardinal location. The sign was randomly given for cardinal and 45° locations. The middle bar (i.e., 0° classification error) represents decoding accuracy plotted in panel (a). (c) The ratio between the number of classification errors away from the nearest cardinal locations (i.e., repulsion) and the number of classification errors toward the nearest cardinal locations (i.e., attraction), separately for the durations of WM encoding (0-200 ms), early WM maintenance (200-850 ms), and late WM maintenance. Classification errors within ±45° range (i.e., orange bars) were used for this analysis because errors outside the range are likely to reflect random noise. The time points were determined based on the stimulus presentation period (0-200 ms), and the first and the second half of the delay period (200-1500 ms). Horizontal black line represents the hypothetical baseline where the numbers of repulsion and attraction errors are identical (i.e., zero bias). Error bars indicate ±1 SEM. *** = $p$ <.001, two-tailed permutation test

I conducted the alpha-based decoding analysis for orientation in Experiment 2. In this experiment, response preparation was not possible and attention-based working memory maintenance was less effective because the location of the target during the 2AFC task was unknown to participants (see Discussion in the main paper). Therefore, I predicted that alpha-based decoding would be weaker compared with ERP-based decoding in this experiment. As can be seen from Figure S2a, alpha-based decoding was significantly greater than chance only for a small fraction of time points (220-880 ms). When it was compared with ERP-based decoding, I found a significant cluster of time points where ERP-based decoding was significantly greater than alpha-based decoding (1 cluster, 80-460, $p <. 001$, two-tailed permutation testing). These results are consistent with the hypothesis that alpha-band activity primarily reflects attention-based support processes whereas sustained ERP activity reflects both the representational contents and attention-based support processes (Bae & Luck, 2018).

Figure S2b shows the signed decoding response error distribution for cardinal/45° orientations and oblique orientations. As in the ERP-based decoding, there were apparent opposite orientation errors, which suggest that participants attended to thick-end of the teardrop on some proportion of trials. More importantly, there were more frequent repulsion errors (i.e., positive errors) than attraction errors (i.e., negative errors) in the decoding error distribution for oblique orientations. When two types of errors were compared via the ratio analysis, I found more repulsion errors for the first and second half of the working memory delay periods but not for the perceptual encoding period (Figure S2c). This result suggests that spatial attention was not categorically biased during stimulus encoding but it became categorically biased during working memory maintenance.

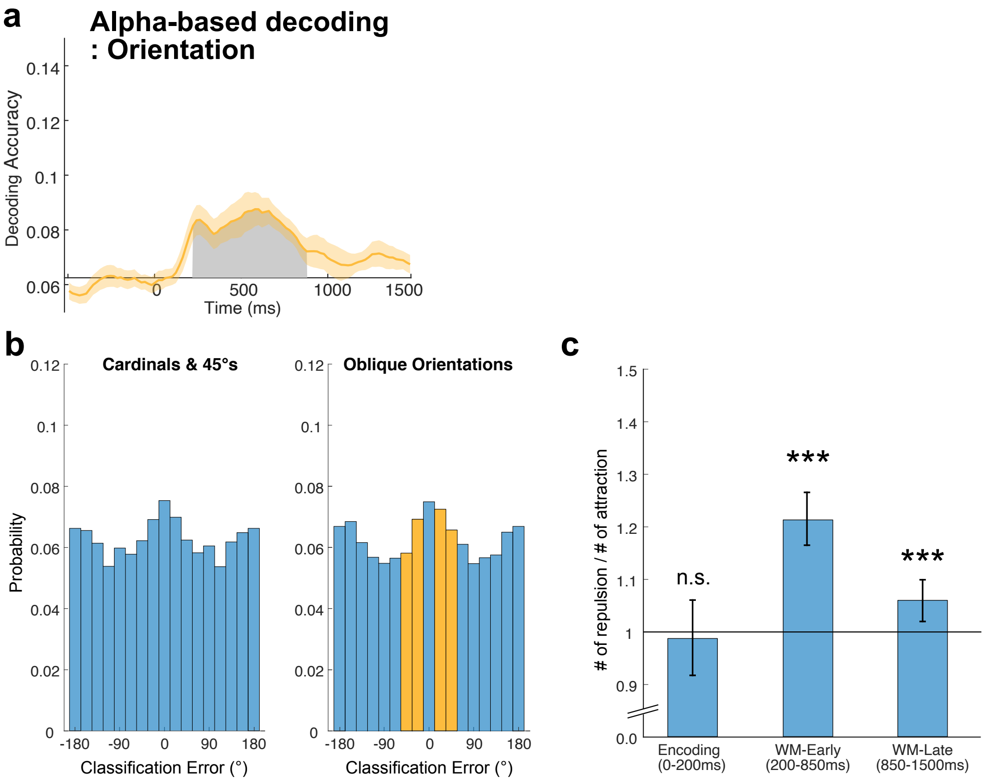


Figure S2. (a) Mean Alpha-based decoding accuracy, averaged across observers in Experiment 2, plotted as in Figure S1(a). (b) Probability distribution of classification errors for cardinal/45° orientations and oblique orientations plotted as in Figure S1(b). (c) The ratio between the number of classification errors away from the nearest cardinal orientation (i.e., repulsion; right two orange bars) and the number of classification errors toward the nearest cardinal orientation (i.e., attraction; left two orange bars), plotted as in Figure S1(c).

2. <u>Decoding without trials with potential eye movements</u>

ICA-based correction for eye movement may not correct for other differences in neural activity that may result from sustained changes in eye position. To ensure that the decoding was not based on signals related to eye position, I conducted an additional set of decoding analyses by excluding trials that could potentially involve systematic shifts in eye position.

I first computed the mean HEOG (Right EOG – Left EOG) and VEOG (Lower EOG – Upper EOG) voltages over epoch, and subtracted the mean pre-stimulus voltage to correct for the baseline voltage offset. I then converted the HEOG and VEOG voltages into a vector (in units of degrees rather than units of $\mu$V) representing the angle and amplitude of the eye position relative to the fixation point, using normative scaling values for HEOG (16 $\mu$V/°) and VEOG (12

$\mu$V/°)(Lins et al., 1993). I excluded trials from decoding analyses if the amplitude of the eye movement for a given trial was greater than 1° (which is less than the half of the radius of the invisible ring) in any direction in Experiment 1 and greater than 0.5° (which is less than the half of the size of the teardrop) in Experiment 2. This procedure excluded approximately 28% of the trials in Experiment 1 and 44% of the trials in Experiment 2. But the excluded trials are likely to include eye movement-irrelevant large voltage fluctuations in a given trial. The exclusion process led both to a smaller number of trials and an unequal number of trials for each stimulus class. Consequently, I found a stimulus class with smallest number of trials and used that number of trials for all other stimulus classes in the decoding analysis and this necessarily decreased signal-to-noise ratio during the averaging process.

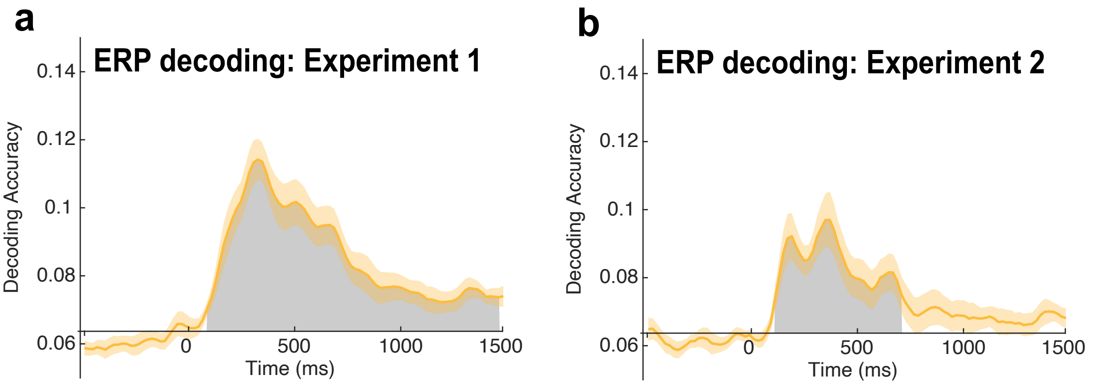


Figure S3. Mean decoding accuracy without trials with large eye movements for (a) Experiment 1 and (b) Experiment 2. The orange shading indicates ±1 SEM. The grey area indicates clusters of time points where the decoding was significantly greater than chance (=1/16) after correction for multiple comparisons.

Despite the reduced number of trials for averaging, the decoding of spatial location in Experiment 1 produced a large cluster of significant time points (Figure S3a, 1 cluster, 80-1500 ms, $p < .001$, one-tailed permutation test). The decoding of orientation in Experiment 2 was noticeably weaker than the main results due to the use of more strict exclusion criteria but still produced above-chance decoding (Figure S3b, 1 cluster, 100-700 ms, $p <.001$, one-tailed permutation test). These results suggest that the main decoding results were not merely driven by systematic eye movements.

3. <u>Categorical biases in decoding when the decoding was done without trials with potential eye movements</u>

As can be seen from Figure S4, decoding without trials with large eye movements exhibited the same pattern of categorical biases in both experiments, demonstrating that the main results were not merely driven by systematic eye movements.

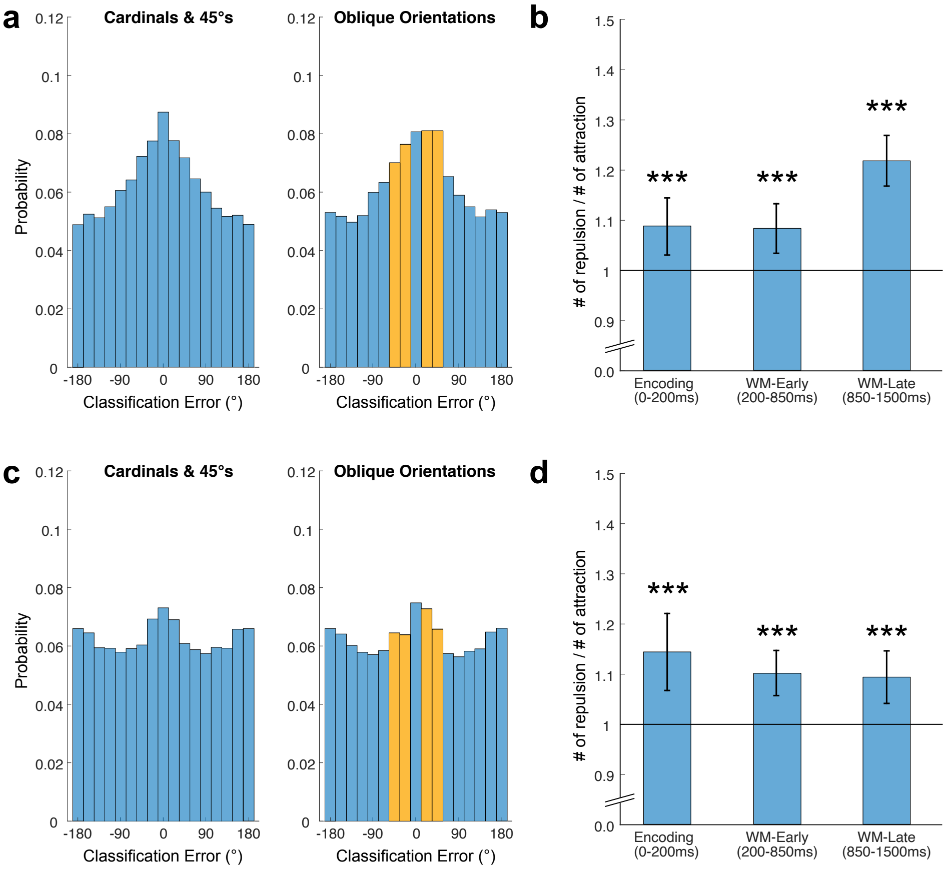


Figure S4**.** (a) Probability distribution of classification errors for cardinal/45° locations and oblique locations from the decoding without trials with large eye movements in Experiment 1 plotted as in Figure S1(b). (b) The ratio between the number of classification errors away from the nearest cardinal location (right two orange bars) and toward the cardinal locations (left two orange bars) in the decoding without trials with large eye movements in Experiment1. (c),(d) Corresponding figures of panels (a) and (b) for the decoding in Experiment 2. *** = $p$ <.001, two-tailed permutation test
